# A simple method of estimating rabies outbreak sizes from phylogenies

**DOI:** 10.64898/2026.06.01.729206

**Authors:** Rowan Durrant, Christina Cobbold, Elaine Ferguson, Kirstyn Brunker, Criselda Bautista, Katie Hampson, Jonathan Dushoff

## Abstract

The ability to estimate the size of a disease outbreak is beneficial both for coordinating an outbreak response and for retrospectively evaluating the effectiveness of response measures and surveillance. Methods such as seroprevalence surveys and epidemiological modelling can be used for this purpose, but are not suitable for every outbreak. We develop a simple formula to estimate the proportion of outbreak cases sequenced, using only the per-generation substitution rate and the length of a phylogenetic tree. This proportion can then be used to estimate the outbreak size. We validate this method using subsampled data from fully observed simulated outbreaks, and then apply it to an ongoing rabies outbreak in the Philippines. We find that this method produces accurate estimates of outbreak size and can be applied to outbreaks with substitution rates and R_0_ values within the estimated range for rabies. We estimated that approximately 19% of cases from the outbreak in the Philippines had been sequenced, suggesting better than average case detection, with a resulting outbreak size estimate consistent with previous analyses. With this method and the associated R package POSER (https://github.com/RowanDurrant/poser), we aim to make rabies outbreak size estimation more accessible, but further work is required to improve its applicability to other diseases.

## Introduction

In the early stages of responding to a disease outbreak it is often difficult to grasp the scale of the problem. Estimates of the outbreak size are key for planning and prioritization, and for evaluating the effectiveness of response measures. Current methods for estimating outbreak sizes or disease prevalence can involve epidemiological modelling (Choi & Ki 2020; Fraser *et al*. 2009; Shutt *et al*. 2017; Zhang *et al*. 2020), randomised testing (Pouwels *et al*. 2021), or serological surveys (Gérardin *et al*. 2008), though these are often impractical, and sometimes impossible.

Rabies is an example of a disease where estimates of outbreak sizes would be useful, but to date have been unobtainable. As with most zoonoses, the majority of rabies infections go either undetected or unreported as dogs in regions experiencing outbreaks are often free-roaming. Because of this, it is difficult to estimate how many rabies cases truly occur in any given outbreak.

As rabies is fatal once the victim becomes symptomatic and infectious, serological surveys cannot be used to estimate historical rabies prevalence, and rabies diagnosis involves acquiring a post-mortem brain sample, which is not practical or appropriate for randomised testing. Furthermore, epidemiological modelling methods often fail to accurately capture the rabies virus’s ability to persist at low prevalence (Mancy *et al*. 2022), and require computational resources and specialised skill sets that may make them inaccessible to local response teams.

Generating genomic data has increasingly become an important part of the outbreak response process, exemplified by the massive testing and sequencing efforts of COVID-19 cases across the world during the pandemic (Ko *et al*. 2022; Meredith *et al*. 2020). Genomic data have the potential to become a useful tool for estimating outbreak sizes. Existing methods leveraging genomic data include Bayesian skygrid (Gill *et al*. 2013; Hill & Baele 2019) and skyline models (Drummond *et al*. 2005; Stadler *et al*. 2013), GlnPipe (Smith *et al*. 2021) and BREATH (Colijn *et al*. 2024). These methods, however, estimate the effective population size (N_e_), an abstract value usually much lower than the real outbreak size (Waples 2022). As such, these methods mainly focus on *relative* growth rates over time (Faria *et al*. 2014). The only method that we are aware of that uses genetic data to explicitly estimate the number of infectious cases is one of joint phylogenetic and epidemiological inference using a particle filtering framework developed by Vaughan *et al*. (2019).

Rabies genomic data have expanded alongside a broader growth in pathogen genomics (Gibson *et al*. 2022; Lushasi *et al*. 2023; Zinsstag *et al*. 2017), with the development of portable sequencing technologies and more cost-effective protocols leading to an increase in capacity for in-country whole-genome sequencing in regions with endemic rabies (Bautista *et al*. 2023; Brunker *et al*. 2020). The rabies virus evolves relatively slowly, however, meaning that even datasets collected over multiple years may lack sufficient temporal signal to support advanced phylodynamic analyses such as skyline or skygrid methods on emerging outbreaks (Durrant *et al*. 2024). Furthermore, local responders are unlikely to have previous experience with these specialised methods, and estimates of N_e_ are likely to be much lower than the true outbreak size. Developing a simple method for outbreak size estimation based on genomic data could aid in making outbreak estimation more accessible in the regions worst affected by rabies.

Phylogenetic trees have certain features that may aid in this goal. When a tip is removed from a phylogenetic tree, two branches are affected: the branch leading to the removed tip is also removed, and its sister branch is merged into the ancestral branch. As the total tree length (the sum of all branch lengths) only reduces by the length of the branch leading to the removed tip but the number of branches reduces by two, the mean branch length will generally increase as tips are removed. This effect is illustrated in Figure 1, where progressively fewer sequenced cases in a hypothetical outbreak produce exponentially shorter trees.

**Figure 1:**
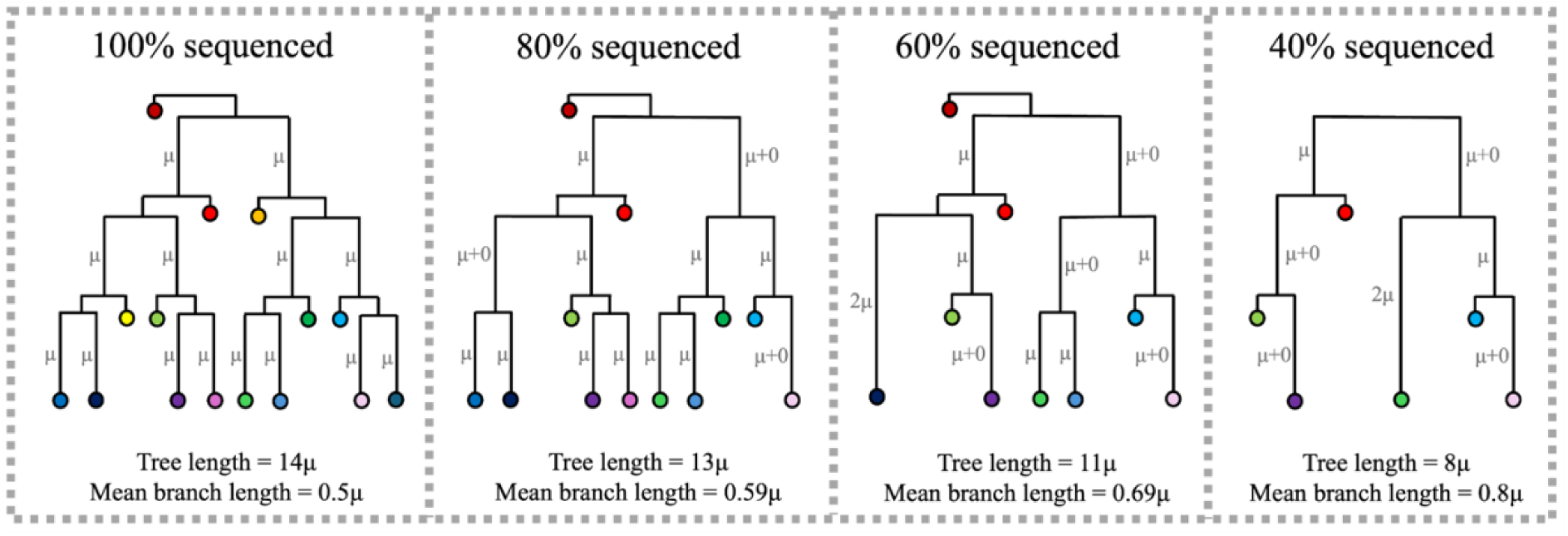
Diagram showing phylogenetic trees of a hypothetical outbreak where every new case gains μ mutations and 100%, 80%, 60% or 40% of cases have been sequenced. Unlabelled branches have a length of 0 (assuming no mutation occurs between onward transmission and sequencing).

We hypothesise that this relationship between the tree length and proportion of cases sequenced, if consistent, could be exploited to estimate the proportion of sequenced cases in an outbreak and thus the outbreak size. In this study we develop a simple method of estimating rabies outbreak sizes using only a phylogenetic tree and an estimate of the per-generation substitution rate, based on observations of phylogenetic tree summary statistics from simulated datasets. We test the accuracy of this method within our simulation framework, explore complications that can decrease its accuracy, and then apply it to data from a recent outbreak of rabies in the Philippines.

## Results

We observed that in our synthetic outbreaks, the tree length (the sum of the branch lengths) had a consistent, approximately square-root relationship with the proportion of sequenced cases (Figure 2). This relationship is consistent across outbreaks of different sizes, and across values for R_0_ and the substitution rate realistic for rabies.

**Figure 2:**
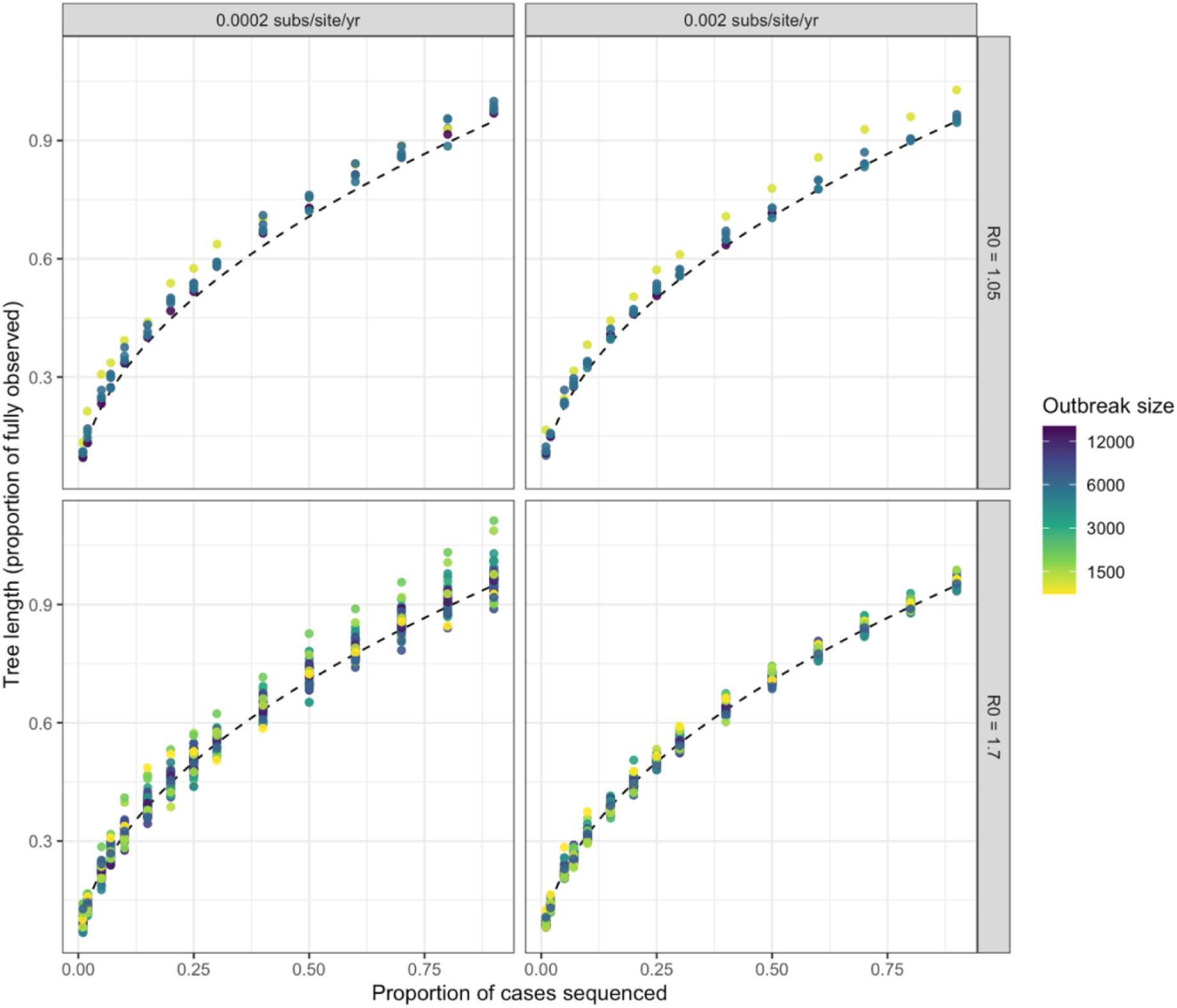
The length of a maximum likelihood tree increases predictably with the proportion of cases in the outbreak that were sequenced. The dashed curve has the formula y = sqrt(x). Point colour represents the total size of the outbreak and is log scaled; facet titles describe the parameters of the simulation.

This pattern suggests the that the following assumption holds:

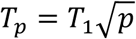

where *T_p_* is the tree length when a proportion p of cases have been sequenced and *T_1_* is the length of the fully observed tree. *T_1_* can be written as

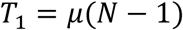

where *μ* is the mean per-generation substitution rate and *N* is the total number of cases in the outbreak, giving:

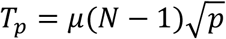

As we know the number of sequences in the tree is *pN*, rewritten as *S*, but not the individual values of *p* or *N*, the above formula can be multiplied through by 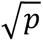 and rearranged to:

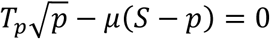

which can be solved for p using the quadratic formula. This allows the value of *p* (the proportion of cases sequenced) to be ascertained and the outbreak size to be calculated.

### Accuracy testing

We tested the accuracy of this formula in simulated rabies outbreaks and found it to be usefully accurate. Over a range of parameter values, we found that the 95th percentile of the absolute value of the log-scaled proportional difference between the estimated and true proportion of cases sequenced was 0.300, where 0 represents perfect estimation. This means that when estimating the size of outbreaks with a true size of 1,000 cases and varying values of *p*, on average 95% of the estimates will fall between approximately 741 and 1,350 cases, assuming that there is no directional estimation bias. When comparing the accuracy of simulations with different values of R_0_, we found that the 95th percentile proportional difference in *p* estimates in outbreaks with an R_0_ of 1.05 is 0.358, whereas the 95th percentile proportional difference when R_0_ is 1.7 is 0.285, indicating that our method is slightly better at estimating *p* for outbreaks with the higher value of R_0_. Our error estimate, the root mean square of the log-scaled proportional difference between the estimated and true proportions of cases sequenced, was 0.142 (0.200 for simulations with R_0_ of 1.05 versus 0.131 for simulations with an R_0_ of 1.7), and this error decreases as the true proportion of cases sequenced increases (Figure 3). Our bias estimate, the mean proportional difference between the estimated and true proportions of cases sequenced, was −0.021 (−0.141 for simulations with R_0_ of 1.05 versus −0.0033 for simulations with an R_0_ of 1.7), meaning that on average the estimated proportions were slightly lower than the true proportion of cases sequenced, but that this bias is stronger in the lower R_0_ simulations.

**Figure 3:**
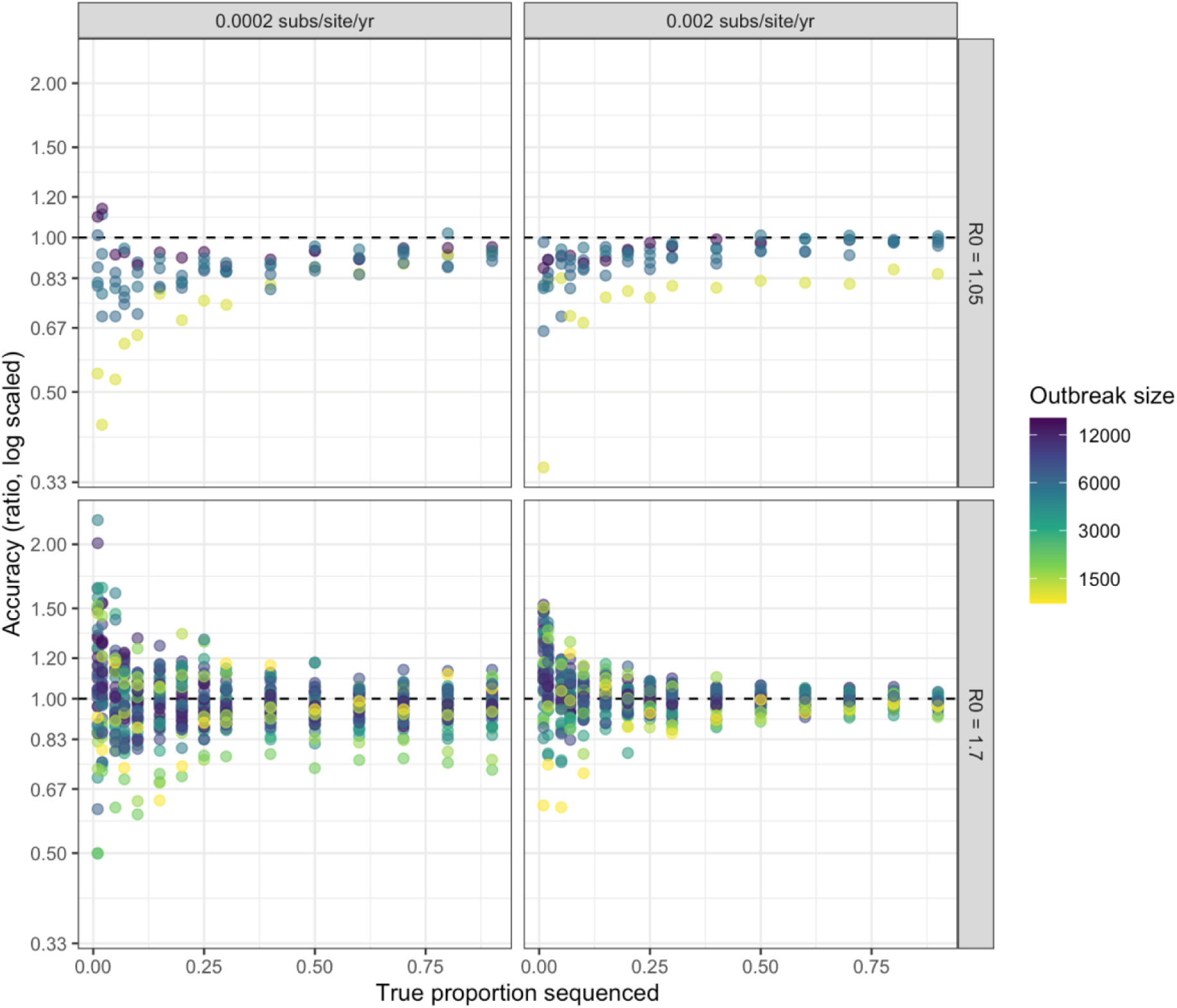
The accuracy of our method in estimating the proportion of cases sequenced when applied to simulated rabies outbreaks with two values of R_0_ and substitution rate. Each point represents a simulated rabies outbreak with at least 500 cases in total. Point colour represents the substitution rate of the simulated outbreak. Facet titles describe the parameters of the simulation. The dashed line represents perfect estimation.

We found that estimates of the proportion of cases sequenced in outbreaks where sequencing was delayed were less accurate (comparing sequencing starting three years versus one year after the index case; Figure 4A). Delays in the start of sequencing resulted in an over-estimation of the proportion of cases sequenced (and therefore an under-estimation of the total outbreak size). This is likely due to some lineages going extinct before sequencing commenced, and these therefore not being represented in the resulting dataset (Figure 4B).

**Figure 4:**
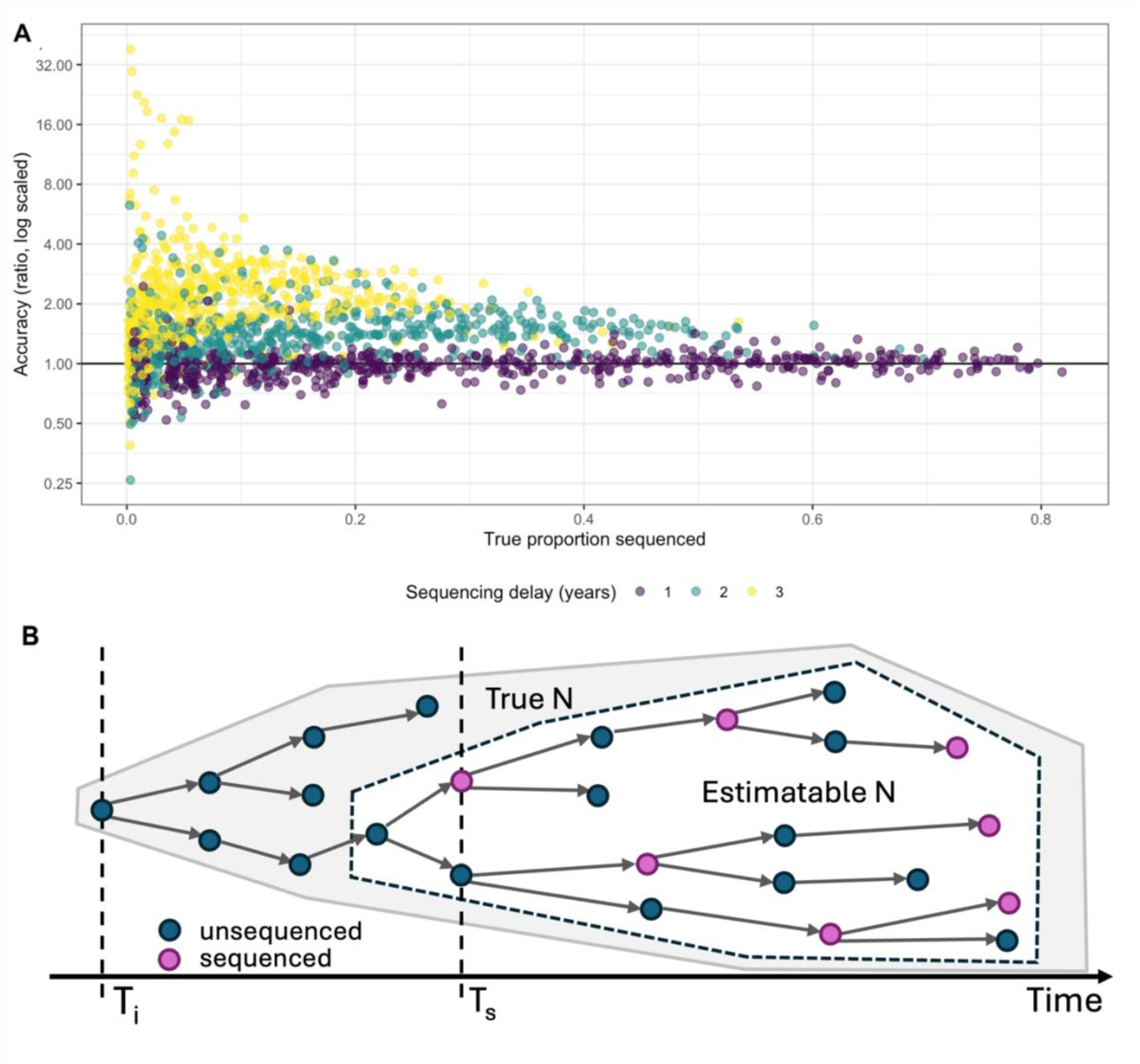
Delays between the start of an outbreak (T_i_) and the initiation of sequencing (T_s_) could lead to the proportion of cases sequenced being overestimated (and therefore the outbreak size underestimated). **(A)** Delays in the start of sequencing efforts can result in reduced outbreak size estimate accuracy. Accuracy is the ratio between the estimated and true proportion of cases sequenced (log scaled), where a value of 1 denotes a perfect estimate, >1 over-estimation, and <1 under-estimation. Point colour represents the delay in the start of sequencing of either one, two or three years into the seven-year outbreak. **(B)** Proposed mechanism by which time delays lead to the underestimation of the outbreak size (N).

Estimate accuracy also varied as the secondary case distribution changed. Holding R_0_ constant at 1.7, we found that estimates for outbreaks simulated with less variance in transmission (a higher negative binomial size parameter) were more accurate than those with a lower size parameter (Figures 5A & B; for reference, our base simulations use a size parameter of 1.33, consistent with observed biting behaviour of rabid dogs; Hampson *et al*. 2009).

**Figure 5:**
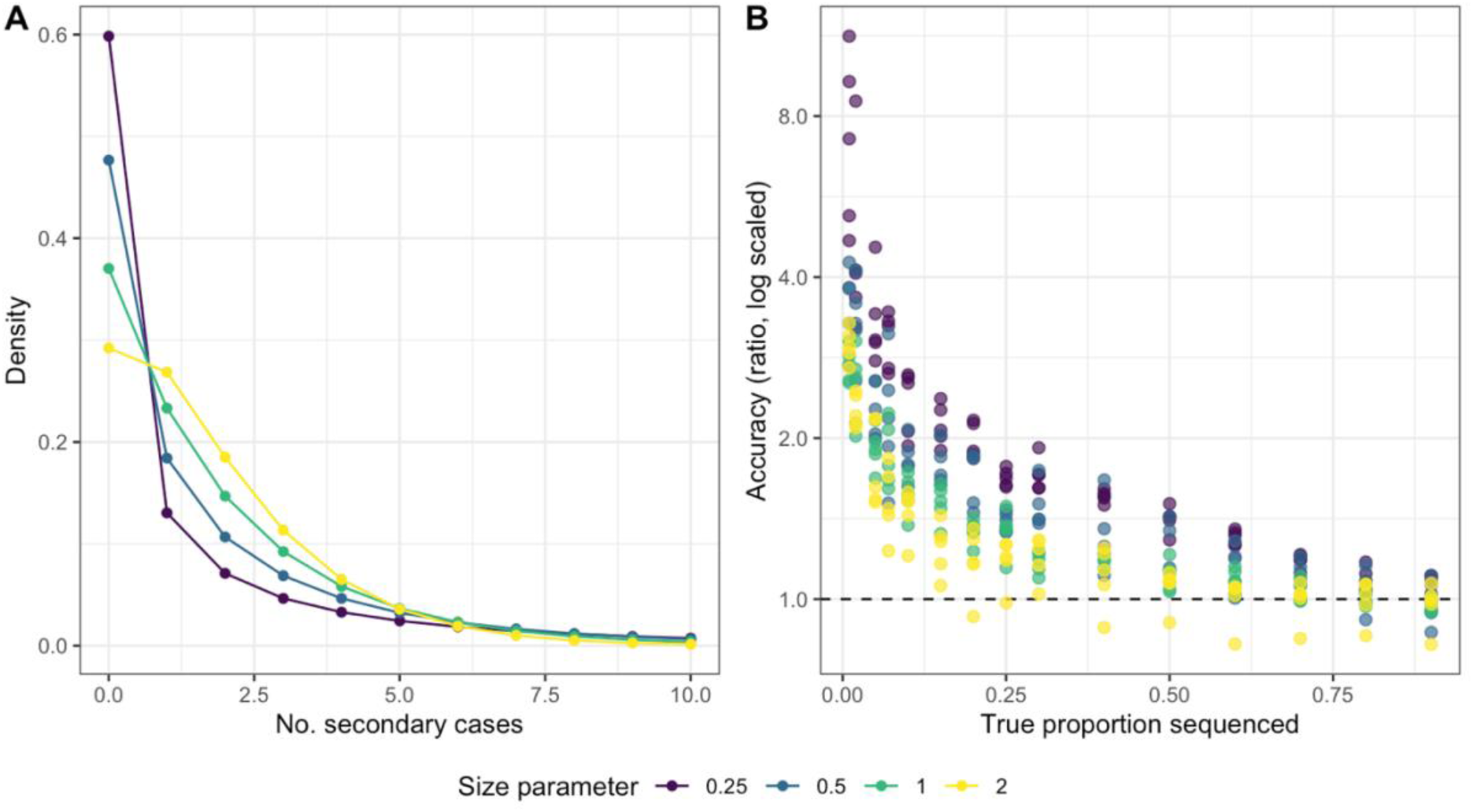
Secondary case distribution affects the accuracy of our method. **(A)** The underlying secondary case distributions investigated. **(B)** As the negative binomial size parameter increases, the accuracy of the outbreak size estimation using our method increases. Point colour represents the size parameter of the negative binomial distribution used to determine the number of secondary cases.

**Figure 6:**
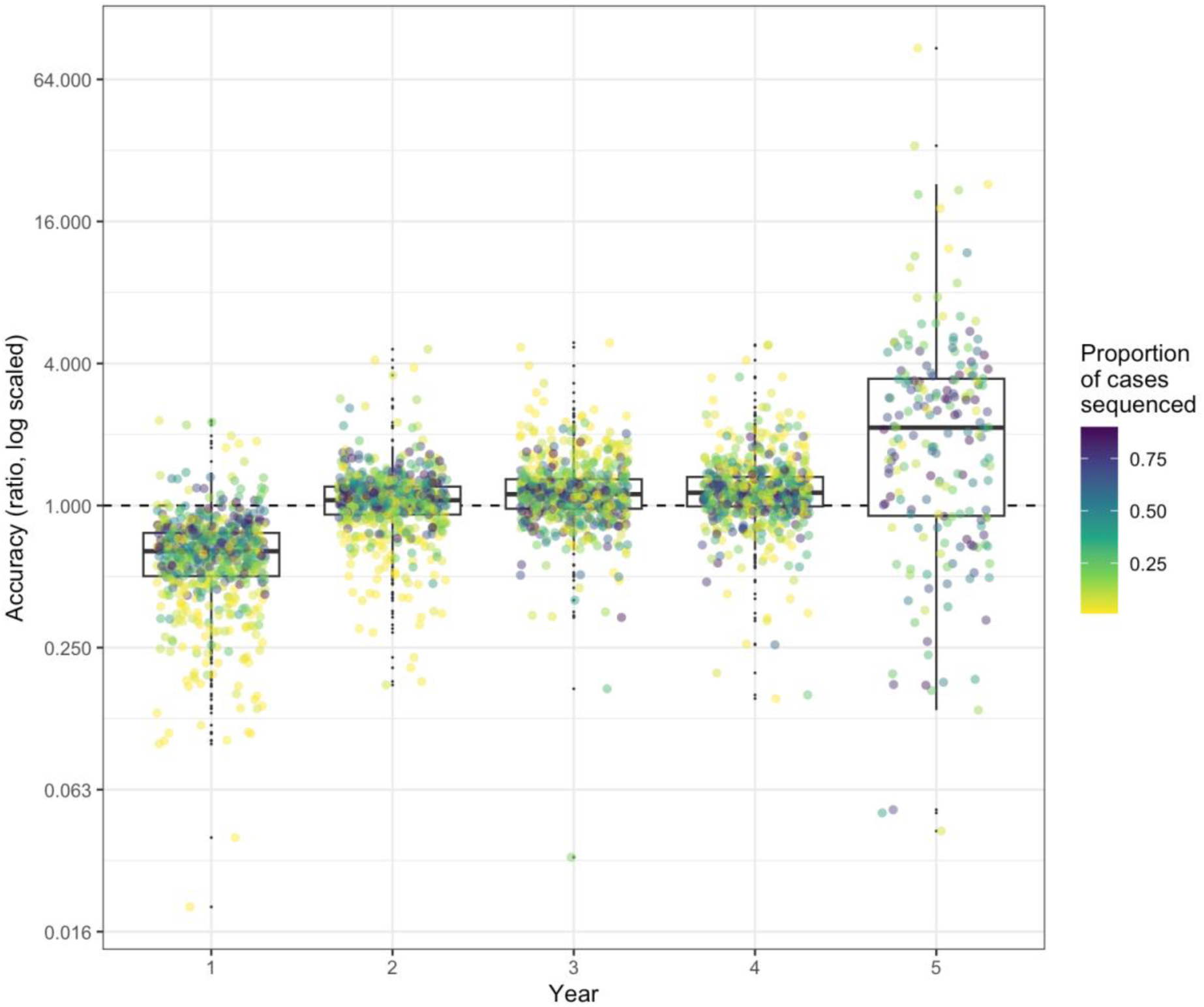
Accuracy in estimating the annual number of cases varies across the course of the outbreak. Point colour represents the proportion of cases sequenced. The dashed line represents perfect estimation (i.e., the estimated annual number of cases is exactly equal to the true annual number of cases).

### Estimating changing prevalence over time

We estimated annual case numbers using our method by sequentially pruning the phylogeny of tips sampled in the latest year (i.e., only including cases that occurred prior to that year) and tested their accuracy using simulations with an R_0_ of 1.7 and a substitution rate of 2×10^−4^ substitutions per site per year. Annual estimates were less accurate than estimates for the full outbreak, (root mean squared proportional difference of 0.581, and 95th percentile proportional difference of 1.25 for the annual estimates, compared to 0.14 and 0.30 respectively for the overall outbreak size estimate) but approximately as biased (bias of −0.020 for annual estimates compared to −0.021 for overall estimates). This suggests that 95% of estimates for years where 1000 cases occurred would be between 287 and 3480 cases without accounting for directional bias, which severely decreases the usefulness of this estimate compared to that of the overall outbreak size. However, this reduction in accuracy appears to be largely driven by the estimates for the first and fifth years of the outbreak. Accuracy for case number estimates in the intermediate years is higher (proportional difference error of 0.362, 95% percentile of 0.74, bias of 0.089) but still worse than the overall outbreak estimate. If reached, the fifth year of these simulations tends to contain very few cases (i.e., <10), and hence fifth year estimates may be based on very small numbers of additional sequences.

### Application to rabies outbreaks

We applied this method to an outbreak of rabies in the Romblon province of the Philippines, incorporating the error observed in the overall synthetic outbreak estimates into a confidence interval. Between September 2022 and March 2023, 29 cases were detected, and 24 of these were sequenced; through phylogenetic analysis it was determined that these cases belonged to distinct genetic clusters that were the result of at least three separate introductions of rabies to the province (Yuson *et al*. 2024). Using the 14 sequences belonging to the largest of these clusters, we estimated that 18.6% (95%CI: 13.9% −24.6%) of cases from this cluster were sequenced, indicating 75 (95%CI: 57 −102) cases occurred in this cluster before March 2023 (Figure 7). Applying this sequencing probability to the full outbreak, we estimate that 129 cases (95%CI: 98 −175) occurred in total during this period, corresponding to 22.5% of circulating cases being detected. Computation time was approximately 0.1 seconds when run locally in R version 4.5.2 on a 2023 MacBook Pro.

**Figure 7:**
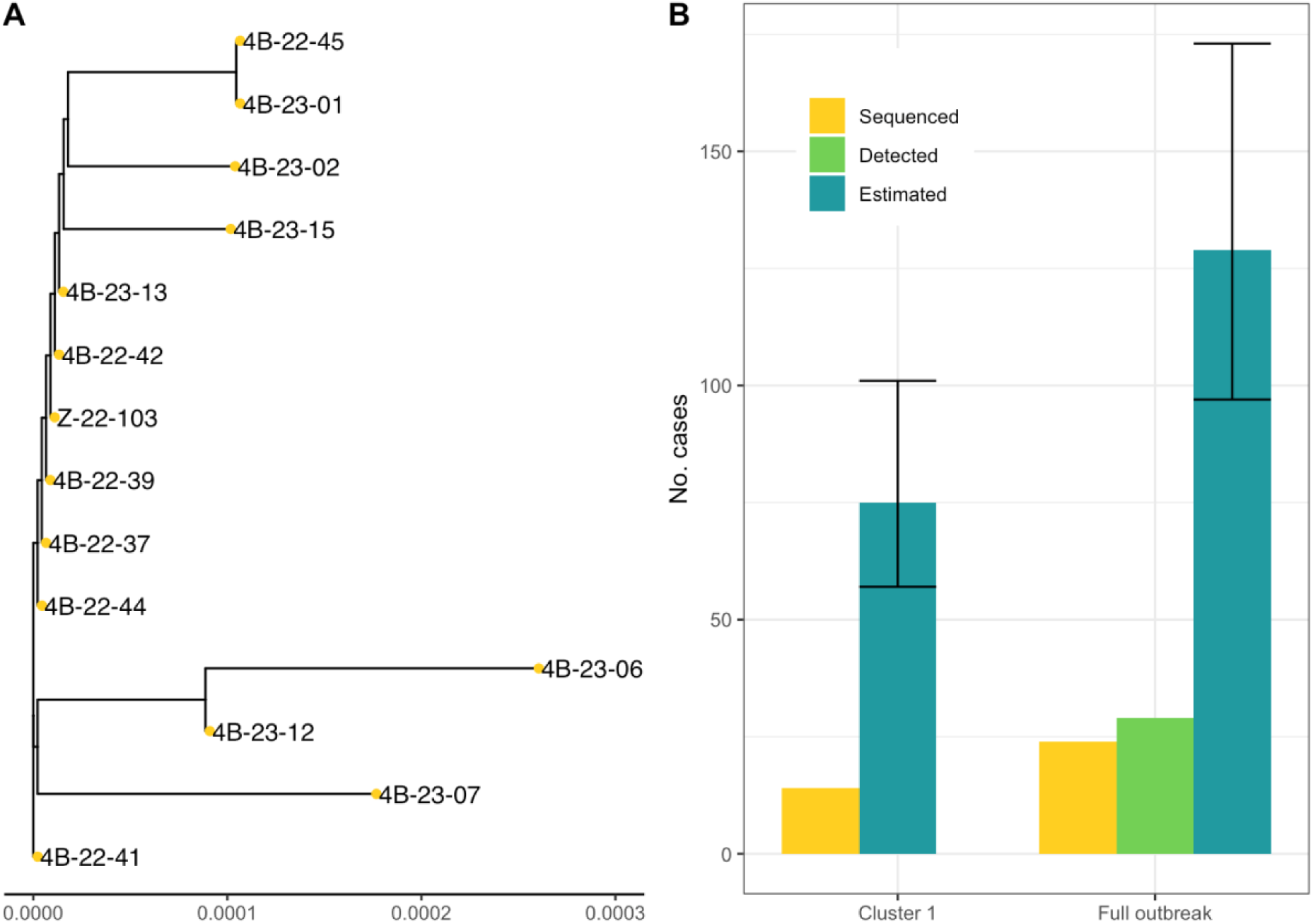
The estimated size of the Romblon rabies outbreak suggests extensive undetected transmission. **(A)** The cluster 1 phylogeny used to estimate the outbreak size; the x-axis is scaled in substitutions per site. **(B)** The numbers of sequenced, estimated and detected cases for the cluster used in our method and the full outbreak, assuming no detection bias between clusters.

## Discussion

Here we describe a novel method of estimating rabies outbreak sizes from phylogenetic trees using minimal computational resources, which, despite its simplicity, has high accuracy. This presents an improvement in accessibility for users in the regions most affected by rabies over existing methods such as modelling and Bayesian phylogenetic methods, which to our knowledge have not yet been successfully applied to rabies outbreaks for the purpose of outbreak size estimation. We estimated that 129 cases (95%CI: 98 −175) occurred in the current Romblon outbreak up to March 2023. When compared to the 29 confirmed cases detected in this period, this suggests considerable undetected transmission occurred in Romblon, consistent with the findings of the original outbreak report (Yuson *et al*. 2024). Despite these levels of undetected transmission, surveillance performance appears to be relatively good compared to many rabies-endemic regions, where case detection is estimated to rarely exceed 10% (Townsend *et al*. 2013).

Our method relies on the assumption that sequencing effort is even over time and unbiased, as delays in sequencing leads to overestimation of the proportion of cases sequenced. In reality, sampling in rabies outbreaks in remote settings is likely to be biased, as sampling usually only commences after a human exposure is reported (Townsend *et al*. 2013), leading to delays, and sampling based on contact tracing may overrepresent cases with descendants. A potential solution to this problem would be to calculate annual estimates as described above, discarding the first annual estimate as this value would represent an estimate of all cases that may have occurred before the end of the first sequencing year, and is likely to be an underestimate due to the unknown existence of lineages that have gone extinct before sequencing began. As the case numbers per interval are calculated from the difference between two cumulative estimates, these intervals do not need to be equal in length; in our example, the few sequences available from year five could be pruned at the same stage as the sequences from year four to form an estimation interval from the start of the fourth year until the end of the outbreak, potentially improving estimate accuracy over this period compared to strictly annual estimates. In very densely sampled outbreaks this interval could be shortened to provide finer resolution case number estimates, or lengthened in sparsely sampled outbreaks.

An important aspect to consider when applying this method is sensitivity to the per-generation substitution rate parameter. In many emerging rabies outbreaks there is insufficient temporal signal to conduct Bayesian phylogenetic analyses on sequences from the outbreak alone (Durrant *et al*. 2024), so using substitution rate values from BEAST log files of a dataset wider than the outbreak being investigated to calculate this value may be tempting. We used a BEAST log file from analysis conducted on a Philippines-wide dataset in our Romblon outbreak size estimates, but this may lead to the results being affected by the time-dependent rate phenomenon. As purifying selection has a strong effect on RNA virus genomes (Hughes & Hughes 2007), and the substitution rate estimate decreases over a wider temporal sampling window because of this (Duchêne *et al*. 2014), the estimated per-generation substitution rate using the wider dataset may be lower than the per-generation substitution rate of the dataset being investigated, leading to an under-estimation of the proportion of cases sequenced. Where Bayesian phylodynamic analyses cannot be conducted, either due to computational costs or lack of temporal signal in the dataset, substitution rate distributions could be generated from informative priors such as scientific literature estimates.

This method relies on the observed tree length having an approximately square-root relationship with the proportion of cases sequenced. Such a relationship has been previously noted in a study that down-sampled a large dataset of human genomes (Karczewski *et al*. 2020). Annual case estimates for simulated outbreaks without temporal sequencing bias underestimated the number of cases in the first year, which may be due to this year lying in the exponential growth phase of the simulated outbreaks; this, along with the difference in accuracy observed between simulated outbreaks with an R_0_ of 1.05 and 1.7 and the differences in estimation accuracy between outbreaks with different secondary case distributions, suggests that this square-root relationship may not be universal between and throughout outbreaks. Further work is required to determine which epidemiological parameters affect the relationship between the tree length and the proportion of cases sequenced to improve estimation accuracy and widen this method’s applicability to other diseases.

In this study we describe a novel, simple method of estimating rabies outbreak sizes from phylogenetic trees, with potential for application to the evaluation of rabies control measures and assessment of surveillance performance. Rabies control efforts have been increasing in response to the World Health Organization and partners setting a global target (Zero by 30) for ending human deaths due to dog-mediated rabies by 2030 (Minghui *et al*. 2018). As these efforts intensify, incursions and outbreaks are to be expected in newly rabies-free regions (Lushasi *et al*. 2023; Rysava *et al*. 2020). Application of our methods to such outbreaks could be valuable for monitoring progress and maintaining surveillance performance. Endemic settings are complicated by the co-circulation of rabies lineages (Bourhy *et al*. 2016; Mancy *et al*. 2022; Zinsstag *et al*. 2017), but the frequent invasion and extinction dynamics that characterize rabies endemicity could be a rich area for further methodological development and application. In settings working towards elimination, we expect to see lineage extinctions and progressive reductions in prevalence that could be measured through our approach, to evidence progress towards the Zero by 30 goal. Beyond rabies, this method has the potential to be developed further and be applied to other diseases.

## Methods

### Rabies simulation

Rabies outbreaks were simulated in R (R Core Team 2025) using a negative-binomial branching process model, as previously described (Durrant *et al*. 2024). Simulations were run for seven years, or until rabies went extinct. Synthetic genetic data were generated for each case in these outbreaks, with a constant time-rate of substitution. All mutations were assumed to occur before onward transmission and sampling, which in this model occur simultaneously, and the generation interval for each case was drawn from a lognormal distribution fit to observed rabies generation intervals (Mancy *et al*. 2022). The number of substitutions was drawn from a Poisson distribution, with the mean given by the substitution rate multiplied by the generation interval. The site of each substitution in the 11922-nucleotide genome (the length of a whole-genome RABV sequence) was selected based on site-specific substitution rates, and the replacement nucleotide was chosen based on transition-transversion rates, both of which were estimated using IQ-TREE (Nguyen *et al*. 2015) from a dataset of 127 whole-genome sequences from the Cosmopolitan AF1b clade, dating from 1981 to 2017, acquired from RABV-GLUE (Campbell *et al*. 2022). Simulated substitutions had no impact on viral fitness or sequencing probability.

Synthetic outbreaks were generated with two values of R_0_ (1.05 and 1.7, both within the plausible range estimated for rabies outbreaks (Li 2019); the number of secondary cases is drawn from a negative binomial distribution with the R_0_ value as the mean and 1.33 as the size parameter) and genetic data was generated with two substitution rates (2×10^−3^ and 2×10^−4^ substitutions per site per year, to represent plausible substitution rates across a range of RNA viruses; 2×10^−4^ substitutions per site per year is the approximate substitution rate for rabies (Layan *et al*. 2021), whereas many other RNA viruses have a substitution rate an order of magnitude higher (Sanjuán 2012)). 273 outbreaks originating from a single introduction were generated for each value of R_0_, but only outbreaks containing over 500 cases were kept for analyses, to ensure that phylogenetic trees could be generated for datasets with the sparsest sampling strategy used. This resulted in 6 outbreaks with an R_0_ of 1.05 and 40 outbreaks with an R_0_ of 1.7, ranging in size from 1,048 to 15,287 cases. Subsets of the resulting synthetic sequences were then randomly sampled to emulate between 1% and 90% of cases being sequenced.

### Accuracy testing

We tested our method using the synthetic data described above. The per-generation substitution rate (substitutions per site per generation; μ) was calculated by multiplying the mean generation interval length (years per generation) of the cases in the subsampled dataset by the substitution rate used in the simulations (substitutions per site per year). Maximum likelihood trees were generated using IQ-TREE, and the tree length was calculated in R using the *ape* package (Paradis & Schliep 2019). We calculate the ratio of the estimated proportion of cases sequenced to the actual proportion of cases sequenced, where a value of 1 represents perfect estimation (i.e., the estimated and true values are equal), and use the natural log of this ratio as our measure of proportional difference.

We then used the accuracy data described above to calculate confidence intervals for our point estimates. We fit a generalised linear mixed model to the accuracy data using the R package *glmmTMB* (Brooks *et al*. 2017), where the log-scaled ratio between the estimated and true values of *p* was the response variable and a B-spline fitted to the estimated value of *p* with four degrees of freedom was the predictor variable. This B-spline is also used as the formula to estimate dispersion. Using the *predict()* function we can extract the expected mean and standard deviation of the ratio between the estimated and true values of *p* for the estimated value of *p*, and fit a normal distribution using these parameters.

We explored the effect of sampling delays on the accuracy of our estimates by imposing a delay (of either one or three years) after the index case was introduced until sampling starts. To investigate the effect of secondary case overdispersion, we adjusted the size parameter of the negative binomial distribution used to determine the number of secondary cases to be either 0.1, 0.5 or 1, maintaining a mean of 1.7. For both the varying temporal bias and secondary case distribution scenarios we used only outbreaks with an R_0_ of 1.7 and a substitution rate of 2×10^−4^ substitutions per site per year.

### Estimating changing prevalence over time

Annual case numbers were estimated by first estimating the full outbreak size using every sequence in the dataset, and then removing the latest year’s tips from the phylogeny using the *drop.tip()* function from the *ape* package, and re-calculating the outbreak size until only the first year of sequence data remains. This gives us estimates of the cumulative outbreak size per year; the difference between these values gives the annual case numbers.

### Application to rabies outbreaks

In September 2022 the first rabies case since 2012 was detected in the Romblon province of the Philippines, which had previously been declared rabies-free. The resulting outbreak response has included enhanced surveillance through integrated bite case management (Swedberg *et al*. 2023), with 45 confirmed cases being detected between September 2022 and September 2023. Two of these cases were human deaths, and the remaining 43 were domestic dogs. From these confirmed cases 24 whole-genome sequences were produced using an Oxford Nanopore MinION-based protocol (Bautista *et al*. 2023), with the latest case to be sequenced being identified on 1st March 2023; 16 of the 45 detected cases occurred after this date (29 cases were detected during the sequencing interval). Further details of the outbreak and the ensuing response, including details of how the RABV sequences were generated, are available in Yuson *et al*. (2024).

A maximum likelihood tree was generated using all 24 available sequences from Romblon using IQ-TREE, and a subtree was extracted consisting of the 14 sequences identified as potentially resulting from a single introduction to the region (“cluster 1”). The remaining sequences were identified as likely resulting from separate introductions, and were too few in number to base estimates on (1, 1, 3 and 5 sequences per cluster). We used 9,002 values from the posterior distribution for substitution rate acquired from the BEAST (Drummond *et al*. 2012) log file of a Philippines-wide tree and the mean generation interval estimated through transmission tree reconstruction (Yuson *et al*. 2024) to calculate the per-generation substitution rate parameter. This tree was generated from 299 whole-genome sequences using a coalescent Bayesian skyline plot with a GTR F+I+G4 uncorrelated relaxed substitution model, with a 10% burn-in removed (Bautista 2025). We then solved our formula for *p* using these parameter values. For each of the 9,002 predictions we then incorporated the error introduced from the method itself as described above, resulting in a distribution of values for each, and then combined the distributions with the *distplyr* package’s mix() function (Coia 2025). We then extracted the mean and 2.5% and 97.5% percentiles from the resulting distribution as the 95% credible interval.

## Data Availability

All code and data used in this study are available in the following GitHub repository: https://github.com/RowanDurrant/branch_lengths. Our R package can be found and installed at https://github.com/RowanDurrant/poser.

## Acknowledgements

The authors would like to thank Ben Bolker, Dan Haydon and Caroline Colijn for their useful suggestions and feedback. This work was supported by the EPSRC DTP (EP/T517896/1 to the University of Glasgow), Wellcome (207569/Z/17/Z, 224520/Z/21/Z to KH), the MRC (MR/X002047/1 to KB) and the Canadian Institutes of Health Research.

